# Recollection-related hippocampal fMRI effects predict longitudinal memory change in healthy older adults

**DOI:** 10.1101/2020.03.19.999060

**Authors:** Mingzhu Hou, Marianne de Chastelaine, Manasi Jayakumar, Brian E. Donley, Michael D. Rugg

**Author notes:** Corresponding author:* Mingzhu Hou, 1600 Viceroy Drive, Suite 800, Dallas, TX 75235.

## Abstract

Prior fMRI studies have reported relationships between memory-related activity in the hippocampus and in-scanner memory performance, but whether such activity is predictive of longitudinal memory change remains unclear. Here, we administered a neuropsychological test battery to a sample of cognitively healthy older adults on three occasions, the second and third sessions occurring one month and three years after the first session. Structural and functional MRI data were acquired between the first two sessions. The fMRI data were derived from an associative recognition procedure and allowed estimation of hippocampal effects associated with both successful associative encoding and successful associative recognition (recollection). Baseline memory performance and memory change were evaluated using memory component scores derived from a principal components analysis of the neuropsychological test scores. Across participants, right hippocampal encoding effects correlated significantly with baseline memory performance after controlling for chronological age. Additionally, both left and right hippocampal associative recognition effects correlated significantly with longitudinal memory change after controlling for age, and the relationship with the left hippocampal effect remained after also controlling for left hippocampal volume. Thus, in cognitively healthy older adults, the magnitude of hippocampal recollection effects appears to be a robust predictor of future memory change.

## 1 Introduction

As they age, healthy adults typically demonstrate reduced performance in multiple cognitive domains. One of these domains is episodic memory (Nyberg and Pudas, 2019) which, following Tulving (1983), we define here as memory for unique events. Numerous functional magnetic resonance imaging (fMRI) studies have investigated the neural correlates of episodic memory processing in older adults (for review, see Rajah et al., 2015; Tromp et al., 2015), but whether any of these correlates are predictive of performance on standardized memory tests, or changes in test performance over time, remains largely unknown. In the present study, we focused on the possible predictive roles of encoding- and retrieval-related neural activity in the hippocampus, a structure that is both necessary for episodic memory (Eichenbaum, 2017; Moscovitch et al., 2017) and has repeatedly been implicated in age-related memory decline (e.g. Persson et al., 2006; Rodrigue et al., 2013; for review, see Leal and Yassa, 2015).

As we discuss below, fMRI studies have employed two classes of experimental contrasts to examine the neural correlates of episodic memory at encoding and retrieval, respectively. The ‘subsequent recollection procedure’ entails contrasts between the neural activity elicited by study items according to whether the items were successfully recollected on a subsequent memory test or were judged as studied merely on the basis of an acontextual sense of familiarity. Resulting differences in neural activity will be referred to below as *encoding effects.* Similarly, the neural correlates of successful recollection – hereafter *recollection effects* –are identified by contrasting the neural activity elicited by memory test items according to whether the items were successfully recollected or judged as studied on the basis of familiarity alone. As is described in more detail in the Materials and Methods, in the present study these contrasts were performed in the context of an associative recognition test in which participants were required to discriminate between pairs of test words presented in the same pairing as at study (‘intact’ pairs), and test pairs comprising words that had been studied on two different study trials (‘rearranged’ pairs). The neural correlates of recollection were operationalized as the contrast between neural activity elicited by intact items according to whether the items were correctly judged intact, or wrongly judged as rearranged (see de Chastelaine et al., 2016a, 2016b, for a detailed rationale for this contrast).

Numerous cross-sectional studies have examined the effects of age on fMRI correlates of memory encoding (e.g. de Chastelaine et al., 2011, 2016a; Logan et al., 2002; Miller et al., 2008; Park et al., 2013; for review, see Maillet and Rajah, 2014) and retrieval (e.g. Daselaar et al., 2003; de Chastelaine et al., 2016b; Duarte et al., 2008; Wang et al., 2016; Wang and Giovanello, 2016; for review, see Giovanello and Dew., 2015). In the case of the hippocampus, findings from both encoding and retrieval studies are mixed; whereas some studies reported null effects of age on encoding- or retrieval-related hippocampal effects (e.g. Angel et al., 2016; Dulas and Duarte., 2016; Miller et al., 2008; Park et al., 2013), others have reported that the effects were either enhanced (e.g. Dulas and Duarte., 2011; Duverne et al., 2008) or attenuated in older adults (e.g. Daselaar et al., 2006; Dennis et al., 2008, Dulas and Duarte., 2014) (it should be noted however that the interpretation of age differences in functional activity in the presence of age differences in performance on the experimental memory task is arguably problematic, e.g. de Chastelaine et al., 2016b; Rugg and Morcom, 2005).

As was just noted, numerous studies have investigated the effects of age on hippocampal functional correlates of encoding and retrieval. However, a substantially smaller number have examined whether such correlates are associated with performance on the experimental memory task. Moreover, only one study has described relationships between these correlates and performance on standardized memory tests, and only two studies have examined whether hippocampal functional correlates might be predictive of longitudinal memory change. Turning first to associations between hippocampal effects and experimental memory performance, the findings are mixed. Four studies reported age-invariant, positive relationships between the magnitude of hippocampal encoding (de Chastelaine et al., 2016a) or retrieval (Daselaar et al., 2006; de Chastelaine et al., 2016b; Wang et al., 2016) effects and memory performance. By contrast, other studies examining such relationships in samples of older adults have reported *negative* relationships for encoding- (Miller at al., 2008) or retrieval-related activity (Carr et al., 2017), or a null relationship (Dulas and Duarte, 2011, 2016). In a similar vein, Daselaar et al. (2015) reported a negative relationship between hippocampal recollection effects and a composite index of memory function derived from a neuropsychological test battery. We defer comment on possible reasons for these disparate findings until the Discussion.

With the exception of Hantke et al. (2013) and Leal et al. (2017), aging studies examining relationships between hippocampal functional activity and longitudinal memory change have employed measures of hippocampal activity estimated either relative to an implicit baseline, or through non-mnemonic contrasts (O’Brien et al., 2010; Persson et al., 2012; Pudas et al., 2013, 2014; Woodard et al., 2010). Thus, it is not possible to classify these studies according to whether hippocampal activity was encoding- or retrieval-related. One study (Woodard et al., 2010) employed a prospective design and reported that a baseline measure of ‘hippocampal’ activity (an amalgam of activity in the hippocampus and parahippocampal cortex) elicited by a contrast between famous and non-famous names contributed significantly to regression models that predicted whether participants’ memory scores would remain stable or decline over the following 18 months. Hantke et al. (2013) described additional analysis of the same data-set and reported that, unlike in the fame judgment task, neural activity differentiating between ‘old’ and ‘new’ recognition memory judgments did not contribute to prediction of future memory decline.

In another two studies employing longitudinal designs, Persson et al. (2012) reported that decline in left hippocampal activity over 6 years was positively associated with longitudinal memory change over a 20 year period. Similarly, O’Brien et al. (2010) reported that longitudinal decline in right hippocampal activity was positively related to memory decline over 2 years. In two final studies (Pudas et al., 2013, 2014), the relationship between hippocampal activity and retrospective memory change was investigated. Pudas et al. (2013) reported that older adults whose memory performance had remained stable over the preceding 15-20 years demonstrated higher levels of hippocampal activity than did adults in whom memory tended to decline over the same period. In a further analysis of the same data-set, Pudas et al. (2014) reported that older adults’ hippocampal activity was positively related to midlife memory performance in addition to performance at the time of scanning.

Together, the above-mentioned longitudinal findings indicate that hippocampal activity can be predictive of individual differences in longitudinal memory change in healthy older adults. As was previously noted, however, the measures of hippocampal activity employed in these studies cannot easily be understood in terms of neural activity directly related to episodic encoding or retrieval operations. One relevant study is that of Leal et al. (2017), who contrasted hippocampal activity elicited by visual scenes according to whether the scenes were confidently recognized or forgotten on a subsequent memory test. The resulting subsequent memory effect in the right hippocampus was unrelated to change in CVLT Long-Delay Free Recall performance over an average follow-up period of 2.7 years. It seems possible, however, that this null finding is a reflection of the marked divergence between the nature of the experimental items (visual scenes) and standardized test materials (words).

Here, we employed the verbal associative recognition procedure described previously to obtain measures of encoding- and recollection-related hippocampal activity in healthy older adults, and examined the relationships between these measures, baseline verbal memory performance and, most saliently, longitudinal (three year) memory change. In addition, we examined whether any such relationships were moderated by hippocampal volume, which has sometimes been reported to be predictive of memory performance and memory change in older adults (e.g. Gorbach et al., 2017; Rosen et al., 2003). At issue is whether recollection-related hippocampal activity is predictive of either baseline memory performance or longitudinal memory change.

## 2 Materials and Methods

In the present report, we describe neuropsychological test data obtained in three test sessions, separated by one month and three years respectively. The data from session 1 have been described previously (de Chastelaine et al., 2015, 2016a, 2016b, 2017; de Chastelaine et al., in press; King et al., 2018), but the data from the succeeding two sessions have not been previously reported. The data pertaining to relationships between the test scores and structural and functional hippocampal measures have also not been reported previously.

### 2.1 Participants

Participants were 67 heathy older adults recruited from the greater Dallas community. They comprised a sub-set of 69 older adults who received the same neuropsychological test battery (see below) on two sessions spaced over a one-month period and who were eligible for structural and functional neuroimaging. The two additional participants were excluded from all analyses (including the PCA conducted on the session 1 neuropsychological test scores, see below), because of abnormal anatomical scans. Intracranial and hippocampal volumetric data from one participant and hippocampal volumetric data from one additional participant were excluded because of low quality structural images. Functional data from 3 participants were excluded because of near-chance performance on the in-scanner memory task (2 participants) or insufficient ‘associative miss’ trials (1 participant; see below for details about the functional MRI session).

A subsample of 55 participants were re-administered the neuropsychological test battery approximately 3 years after test sessions 1 and 2 [12 older adults did not participate in the session 3 due to death (*N*=1), relocation from the Dallas area (*N* = 5), loss of contact (*N* = 5) or failure to attend *(N* = 1)]. Out of these 55 participants, volumetric data from 1 participant, and functional data from 2 other participants were excluded for one of the reasons mentioned above.

All participants were right-handed, fluent in English by age 5, had no history of neurological or psychiatric disease and had normal or corrected to normal vision. They each gave informed consent according to procedures approved by the UT Dallas and University of Texas Southwestern Institutional Review Boards. They were compensated at the rate of $30 per hour for their participation.

### 2.2 Neuropsychological test battery

The neuropsychological test battery comprised the California Verbal Learning Test-II (CVLT; Delis et al., 2000), the Logical Memory test of Wechsler Memory Scale (WMS-IV), the Digit Span Forward and Backward test of the Wechsler Adult Intelligence Scale Revised (WAIS-R) (Wechsler, 2001), the Digit/Symbol Coding test of the WAIS-R (SDMT), Trail Making Tests A and B, letter and category fluency tests, the Wechsler Test of Adult Reading (WTAR; Wechsler, 2001) and Raven’s Progressive Matrices (short version) (see Table 2).

Following the initial administration of the test battery, potential participants were excluded from the MRI session if they had 1) scores > 1.5 SDs below the age-appropriate norm on any long-term memory sub-test (CVLT or Logical memory) or on any two other tests; 2) an estimated full-scale IQ < 100 as indexed by performance on the WTAR, or 3) a score on the Mini-Mental State Examination (MMSE) < 27.

Participants who met the inclusion criteria were re-administered the test battery approximately one month later (session 2, range = 14-64 days, mean = 32 days). The majority of the participants were also re-tested after approximately 3 years (session 3, range = 2.9-3.2 years, mean = 3.0 years). Session 2 was included in an effort to attenuate possible re-test effects at session 3, which would lead to an underestimation of cognitive change. This approach was based on evidence that re-test effects tend to be greater for an initial re-test session than for subsequent sessions (Salthouse and Tucker, 2008) and, in addition, are evident across inter-test delays of several years (Salthouse, 2009). In the present case, we re-administered the neuropsychological test battery after a period (1 month) too short for the scores to be affected by age-related cognitive change. As is detailed below, the mean of the scores obtained in the two sessions served as the baseline for the assessment of change at session 3. Averaging scores across sessions 1 and 2 not only had the benefit of attenuating session 3 re-test effects, but also of providing more reliable estimates of baseline performance than those provided from a single test session (note however that the results reported below for the relationships between fMRI measures and memory performance and change were essentially identical when session 2 scores were employed as the baseline, see Supplementary Material). Missing values from one participant for SDMT, Trail A and Trail B tests at session 3 were replaced by the mean performance of the remaining participants for that session.

A principal component analysis (PCA) was used to reduce the raw scores from the neuropsychological test battery to scores on latent cognitive constructs (component scores). PCA was conducted on the session 1 test data of the 67 eligible participants who provided scores for that session (see the section of Participants above). The raw scores were standardized prior to being subjected to PCA. The four principal components with eigenvalues > 1 were retained and subjected to Varimax rotation (Kaiser, 1958). The resulting component loadings are given in Supplemental Table 1, where it can be seen that the components can be broadly characterized as representing constructs associated with memory, fluency, speed, and crystallized IQ. To maintain comparability of component scores across sessions, for each test in the full group, we combined the test scores from session 1 and session 2 into the same dataset and standardized them together. The component loadings were then applied to the standardized test scores from each session to obtain the component scores for that session. A similar procedure was used to calculate the standardized component scores for the longitudinal subgroup, with the exception that for each test, the scores from all three sessions were combined into a single dataset and then standardized. In both cases, memory component scores averaged across sessions1 and 2 comprised the baseline scores.

**Table 1.** Demographic information, summary measures of hippocampal volume and intracranial volume for the study participants (standard deviations in parentheses).

| Variable |  |
| --- | --- |
| *Full Group* |  |
| N | 67 |
| Age at Session 1 (yrs) |  |
| *M* | 68.2 (3.6) |
| *Range* | 63 – 76 |
| Gender | 37 F, 30 M |
| Education (yrs) | 17.2 (2.3) |
| Left hippocampal volume (cc) | 3.15 (.43) |
| Right hippocampal volume (cc) | 3.33 (.42) |
| ICV (cc) | 1469.98 (122.22) |
| *Longitudinal subgroup* |  |
| N | 55 |
| Age at Session 1 (yrs) |  |
| *M* | 68.3 (3.7) |
| *Range* | 63 – 76 |
| Gender | 28 F, 27 M |
| Education (yrs) | 17.3 (2.4) |
| Left hippocampal volume(cc) | 3.18 (.41) |
| Right hippocampal volume (cc) | 3.39 (.39) |
| ICV (cc) | 1479.05 (127.32) |

### 2.3 In-scanner associative memory task

The MRI scanning session, during which both functional and structural data were acquired, occurred between the initial two administrations of the neuropsychological test battery (average of 22 days after Session 1). The fMRI procedure has been described in detail previously (de Chastelaine et al., 2016a, 2016b). Briefly, during an initial functional scan, participants encoded a series of 240 trial-unique word pairs in the context of a relational task (which of the denoted objects would ‘fit’ into which) presented in two consecutive study blocks. After the encoding phase, participants exited the scanner and rested. They re-entered the scanner 15 min later for a scanned associative recognition test that was administrated in three consecutive blocks. The test items comprised 160 ‘intact’ word pairs (pairs re-presented from study), 80 ‘rearranged’ pairs (comprising studied words that were re-paired between study and test), and 80 ‘new’ pairs (pairs of unstudied words). Instructions were to discriminate between the three classes of word pair, signaling the judgment on each trial by pressing one of three buttons. For each of the study and test blocks, there were two buffer pairs at the start and two buffer pairs in the middle, which followed a halfway 30-s break. The study pairs were intermixed with 80 null trials and the test pairs were intermixed with 106 null trials.

### 2.4 MRI acquisition

Functional and structural images were acquired with a Philips Achieva 3T MR scanner (Philips Medical System, Andover, MA USA) equipped with a 32-channel head coil. Functional scans were acquired during both the study and test phases. The functional data were obtained using a T2*-weighted EPI sequence incorporating the following parameters: TR = 2 s, TE = 30 ms, flip angle = 70°, FOV = 240 × 240, matrix size = 80 × 78. Each EPI volume included 33 × 3 mm thick slices with a 1 mm inter-slice gap and an in-plane resolution of 3×3 mm. Slices were acquired oriented parallel to the AC-PC line in ascending order and positioned for full coverage of the cerebrum and most of the cerebellum. Following the second functional scanning session, diffusion tensor images (DTI) and high-resolution T1-weighted images were acquired. The T1-weighted images were acquired with an MP-RAGE pulse sequence (TR = 8.1 ms, TE = 3.7 ms, FOV = 256 × 224, voxel size = 1×1×1 mm, 160 slices, sagittal acquisition).

### 2.5 Data preprocessing and analysis

MRI data were preprocessed in SPM8 (Wellcome Department of Cognitive Neurology, London, UK). The functional images were motion and slice-time corrected, realigned and spatially normalized using a sample-specific template generated across young, middle-aged and older adults. The template was created by first normalizing the mean volume of each participant’s functional time series (separately for study and test) with reference to a standard EPI template based on the MNI reference brain (Cocosco et al., 1997; see also de Chastelaine et al., 2015, 2016a, 2016b, 2017; King et al., 2018). The normalized mean images were averaged within each group and the resulting 3 mean images were then averaged to generate a template that was equally weighted with respect to the 3 age groups. Images were resampled into 3 mm isotropic voxels and smoothed with an isotropic 8 mm full-width half-maximum Gaussian kernel. For the purposes of template formation and anatomical localization of functional effects, the T1 images were normalized with a procedure analogous to that applied to the functional image but using as an initial template the standard T1-weighed MNI reference brain.

Given our previous findings indicating that both encoding- and recollection-related hippocampal effects were localized primarily to the anterior hippocampus (de Chastelaine et al., 2016a, 2016b), we elected to quantify these functional effects using the anatomically defined anterior hippocampus as the region of interest (ROI). This approach ensured that we sampled encoding- and retrieval-related activity in an unbiased manner from the same hippocampal voxel sets. Following Poppenk et al. (2013), the hippocampal ROIs were defined as the portions of left and right hippocampus anterior to y = −21 in MNI space. The ROIs were manually traced on the across-group average T1 anatomical template following the hippocampal segmentation protocol used by Arnold et al. (2015) (see below).

There were two events of interest for the analysis of hippocampal encoding effects: intact study pairs that were later endorsed as intact (subsequent associative hits) and intact pairs that were later incorrectly identified as rearranged (subsequent associative misses). Intact pairs later incorrectly identified as new were separately modeled, along with all other study pairs and buffer pairs modeled as events of no interest. Analysis of the fMRI recollection effects adopted a similar approach, but with the events of interest comprising correctly endorsed intact pairs (associative hits) and intact pairs incorrectly identified as rearranged (associative misses). Pairs correctly endorsed as rearranged, new pairs correctly endorsed as new, and intact pairs incorrectly endorsed as new were also separately modeled. As for the encoding data, all other test and buffer pairs were modeled as events of no interest. For both the encoding and retrieval data, the rest breaks were also modeled, along with 6 regressors representing motion-related variance and constants representing means across each scan session. Null trials and interstimulus intervals were implicitly modeled as the baseline.

For each participant, parameter estimates extracted from voxels falling within the anatomically defined hippocampal ROIs were averaged for each event of interest. Encoding effects were operationalized as greater BOLD activity for items that went on to be classed as associative hits than for items that became associative misses. Recollection effects were operationalized as greater BOLD activity for associative hits than for associative misses.

### 2.6 Manual tracing of the hippocampus and estimation of hippocampal volume

Manual hippocampal tracing was performed using 3DSlicer/v.4.4.0 (https://www.slicer.org) on each participant’s T1-weighted images. Following the hippocampal segmentation protocol by Arnold et al. (2015), hippocampal boundaries were defined laterally and medially by the lateral ventricle, anteriorly by the hippocampal-amygdala transitional zone, posteriorly by the crus of the fornix, inferiorly by the subiculum, and superiorly by the alveus. The volume of interest (VOI) included CA1, CA2/3, dentate gyrus/CA4, alveus and fimbria, avoiding subiculum, the entorhinal cortex and the hippocampus-amygdala transitional zone. Hippocampal volume was estimated by summing the number of voxels within the traced regions. Prior to analysis, hippocampal volume estimates were residualized against intracranial volume (ICV), which was traced from every 12^th^ slice of the transverse plane and estimated using Analyze 11 (https://analyzedirect.com/).

### 2.7 Statistical analyses

We were interested in examining the extent to which the encoding effects and hippocampal recollection effects predicted, 1) in-scanner recollection performance, and 2) baseline memory and longitudinal memory change, as indexed by the memory component scores derived from the neuropsychological test data.

Recollection performance (pR) was indexed by performance on the in-scanner associative recognition task and was estimated as the difference between the proportion of correctly endorsed intact pairs (associative hits) and the proportion of intact pairs incorrectly identified as rearranged (associative misses) (de Chastelaine et al., 2016a, 2016b). In the case of the neuropsychological test data, the average of session 1 and session 2 standardized memory component scores (see the section of Neuropsychological test battery above) provided the baseline against which the scores for session 3 were compared.

To examine whether the hippocampal encoding effects were reliable at the group level, and to examine their lateralization, we conducted a 2 (condition: subsequent associative hit vs. subsequent associative miss) × 2 (hemisphere) ANOVA on parameter estimates extracted from the hippocampal ROIs. An analogous 2 (condition: associative hit vs. associative miss) × 2 (hemisphere) ANOVA was conducted on the parameter estimates extracted from the hippocampal ROIs at retrieval.

We used partial correlation analyses to examine whether encoding- and retrieval-related hippocampal effects, the predictors of primary interest, were related either to in-scanner recollection performance measured by pR, or the baseline memory component scores derived from the test battery (see *Neuropsychological test battery* above). In addition, we constructed a series of linear mixed effects models to examine whether the effects were predictive of mean memory performance (averaged across baseline and session 3) or longitudinal memory change. Each model included a random intercept term to accommodate individual differences in baseline memory scores. We included chronological age as a predictor in all these analyses because preliminary analyses indicated that this variable was correlated with both hippocampal functional effects and memory performance with small-to-medium effect sizes (absolute *r*s ranging from .10 to .32). The linear mixed models took the following general form:

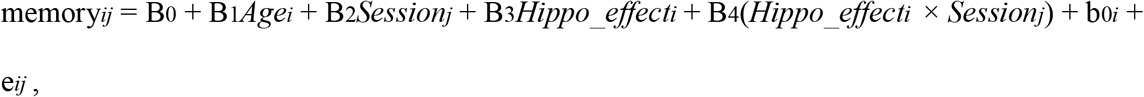

where memory_*ij*_ refers to individual i’s memory performance at session j. Age is participant’s age at baseline, and session is test session (baseline coded as 0, session 3 coded as 1). Hippo_effect refers to either the hippocampal encoding or recollection effect at baseline, and hippo_effect × session refers to the interaction between the hippocampal effect and test session. B denotes fixed-effects estimates, b0 denotes estimates for participant-specific random-effects (i.e. baseline memory scores), and e is the residual error. For those models in which functional activity was found to be a significant predictor, the model was expanded to include hippocampal volume. We constructed additional models to directly examine whether hippocampal volume was predictive of memory performance or memory change.

## 3 Results

### 3.1 Participant characteristics

Demographic information, and summary measures of hippocampal volume and intracranial volume (ICV) for both the full group the longitudinal subgroup are given in Table 1. Consistent with the impression given by the table, the volume of the left hippocampus was significantly smaller than that of the right hippocampus; for the full group, *t*(65) = 6.06, *p* < .001, for the longitudinal subgroup, *t*(53) = 6.62,*p* < .001. Similar findings have been reported previously (Pedraza et al., 2004).

### 3.2 Neuropsychological test performance

Mean performance on each neuropsychological test is given in Table 2 for each of the test sessions. As is evident from the table, for most tests, performance for the first two sessions was well matched between the full group and the longitudinal subgroup. In both groups, there was an overall improvement across tests from session 1 to session 2. In the longitudinal group, mean performance generally showed modest evidence of change between sessions 2 and 3.

**Table 2.** Performance and performance change across sessions for each of the neuropsychological tests (*N* = 67 for full group, *N* = 55 for longitudinal subgroup, standard deviations in parentheses).

| Task | Session |  |  |  |  |
| --- | --- | --- | --- | --- | --- |
|  | 1 | 2 | 1 | 2 | 3 |
|  | Full group |  | Longitudinal subgroup |  |  |
| FAS<sup>a, b</sup> | 45.21 (12.53) | 49.09 (12.74) | 45.04 (12.63) | 48.40 (11.58) | 47.56 (13.26) |
| Logical Memory Composite<sup>a, b, c</sup> | 27.59 (5.39) | 31.73 (5.45) | 27.51 (5.57) | 31.93 (5.39) | 28.39 (5.57) |
| SDMT<sup>a, b, c</sup> | 49.46 (8.50) | 51.91 (8.20) | 49.45 (9.18) | 51.80 (8.73) | 49.09 (8.46) |
| Trail A (ms) | 32.69 (11.24) | 30.25 (11.10) | 31.96 (9.40) | 30.75 (11.71) | 31.89 (10.18) |
| Trail B (ms)<sup>a, b, c</sup> | 75.01 (45.70) | 59.82 (18.51) | 71.44 (30.60) | 59.44 (18.32) | 69.69 (30.54) |
| Digit Span | 18.27 (4.36) | 17.87 (4.24) | 18.25 (4.30) | 17.71 (4.09) | 18.15 (4.26) |
| Category Fluency (Animals)<sup>a, b</sup> | 22.45 (5.56) | 23.96 (5.39) | 22.35 (5.50) | 23.69 (5.44) | 22.93 (5.80) |
| WTAR (Full-Scale IQ)<sup>c</sup> | 112.64 (5.43) | 113.00 (5.16) | 112.95 (5.21) | 113.04 (5.15) | 112.11 (4.63) |
| Raven's | 9.57 (2.13) | 9.91 (1.87) | 9.49 (2.25) | 9.85 (1.87) | 9.56 (2.63) |
| CVLT Hits<sup>a, b</sup> | 14.82 (1.31) | 15.42 (.96) | 14.84 (1.33) | 15.36 (1.01) | 15.25 (1.13) |
| CVLT False Alarms | 1.97 (2.18) | 1.84 (2.53) | 1.93 (2.13) | 1.93 (2.71) | 2.09 (2.52) |
| CVLT Composite<sup>a, b, c</sup> | 12.00 (2.47) | 13.63 (2.02) | 11.95 (2.57) | 13.54 (2.17) | 12.80 (2.55) |

*Note.* a: session 1 ≠ session 2, $p < .05$ for full group; b: session 1 ≠ session 2, for longitudinal subgroup, $p < .05$ ; c: session 2 ≠ session 3, $p < .05$ .

Memory component scores for each test session, and the baseline score averaged across sessions 1 and 2 are shown in Table 3. Performance on session 2 was significantly higher than that on session 1 for both the full group, *t*(66) = 9.69, *p* < .001 and the longitudinal subgroup, *t*(54) = 8.36,*p* < .001. pR (associative recognition performance) for the full group (*M* = .30, *SD* = .15) was closely similar to that for the longitudinal subgroup (*M* = .28, *SD* = .14). Together with the findings for the neuropsychological test battery (see Table 2), this finding provides reassurance that attrition of the sample between sessions 2 and 3 was non-selective in respect of baseline cognitive performance. Baseline memory component scores were significantly correlated with memory scores at session 3, *r* = .87,*p* < .001, indicating high test-retest reliability between baseline and session 3. Finally, pR was moderately correlated with baseline memory component scores in both groups (for full group, *r* = .40, *p* = .001; for longitudinal subgroup, *r* = .48, *p* < .001).

**Table 3.** Standardized memory component score for each session and change score over three years (standard deviations in parentheses).

|  | Session |  |  |  |  |
| --- | --- | --- | --- | --- | --- |
|  | 1 | 2 | 3 | baseline (1&2) | change (1&2 – 3) |
| Full group | -.85 (2.64) | .85 (2.23) | N/A | N/A | N/A |
| Longitudinal subgroup | -.73 (2.69) | .86 (2.30) | -.12 (2.73) | .06 (2.41) | .19 (1.35) |

*Note.* For the longitudinal subgroup, memory scores were significantly higher for session 2 than session 3, $t(54) = 5.37, p < .001$ .

The difference score between baseline and session 3 is also shown in Table 3. A *t* test comparing these scores revealed no evidence for longitudinal memory change at the level of the whole sample [*t*(54) = 1.03, *p* = .308]. Individual memory changes from baseline to session 3 are illustrated in Figure 1. As is evident from the figure, most participants demonstrated relatively small changes in memory over the three year follow-up interval.

**Figure 1.**
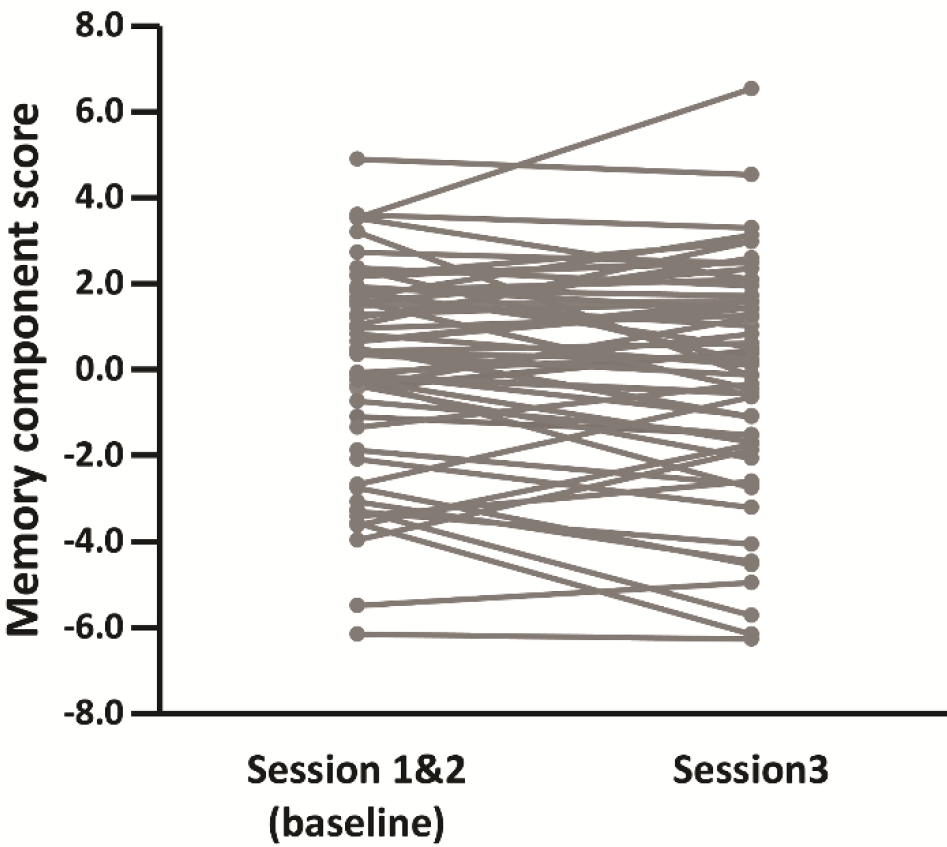
Individual memory component scores at baseline and session 3 for the longitudinal subgroup (*N* = 55). Each line represents memory change for an individual participant.

### 3.3 Functional hippocampal effects

Note that for all analyses of baseline data the findings for the full group and the longitudinal subgroup were equivalent. Therefore, we only report the findings from the full group here. Findings for the longitudinal subgroup can be found in the Supplementary Material.

We first examined whether the functional effects were reliable at the group level, and whether there was any evidence of lateralization in the effects (see Materials and Methods). For the encoding data, neither the main effect of condition nor the condition × hemisphere interaction was significant, *p*s > .075, partial *η*^2^s < .05. An analogous ANOVA conducted on the parameter estimates for associative hits and misses at retrieval revealed a significant effect of condition, *F*(1, 63) = 32.76, *p* < .001, partial *η*^2^ = .34, indicative of greater hippocampal activity for associative hits (*M* = .10) than for associative misses (*M* = -.39). This effect interacted significantly with hemisphere, *F*(1, 63) = 6.38, *p* = .014, partial *η*^2^ = .09, reflecting larger recollection effects in the left hemisphere (*M* = .59) compared to the right hemisphere (*M* = .39). Simple effects analyses indicated that recollection effects were reliable in both hemispheres (*p*s < .001).

### 3.4 Correlations between functional and structural measures and memory performance

Partial correlations (controlling for age) between hippocampal encoding effects, hippocampal recollection effects and in-scanner recollection performance (pR) are given in the left panel of Table 4, while correlations with the baseline memory scores are shown in the right panel of the table. As is evident from the table, with only one exception, the correlations with pR were significant (see also Figure 2). In contrast to the findings for pR, only the right hippocampal encoding effect was significantly correlated with baseline memory scores (see also Figure 3).

**Figure 2.**
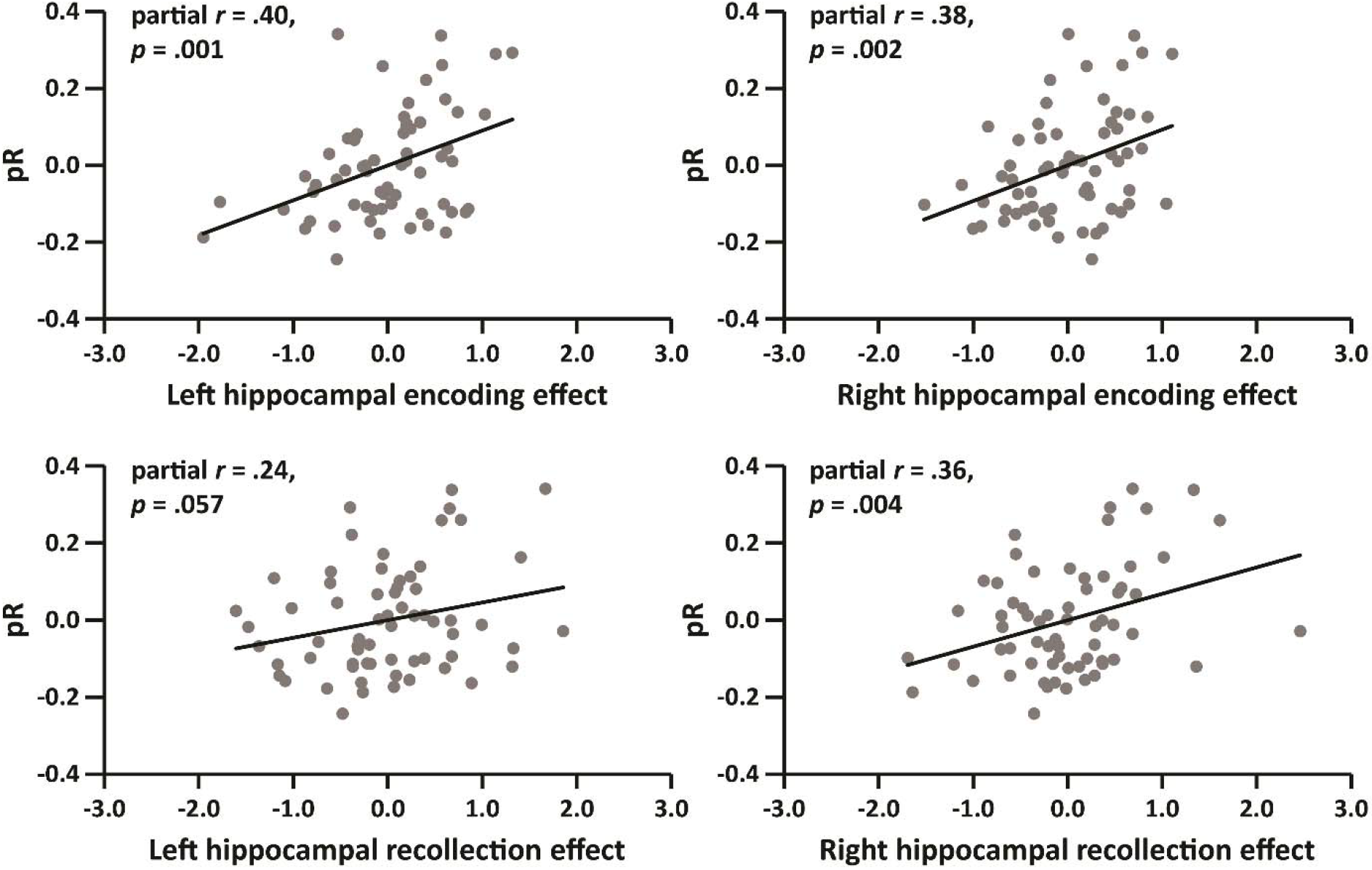
Partial correlations (controlling for age) between pR and functional hippocampal effects (*N* = 64).

**Figure 3.**
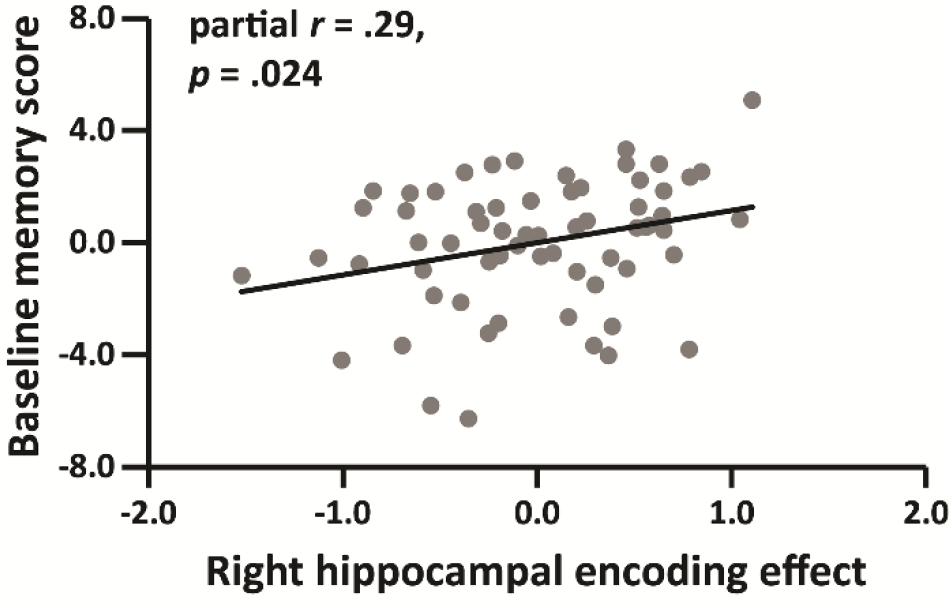
Relationship between the right hippocampal encoding effect and baseline memory score, after controlling for age (*N* = 64).

**Table 4.** Correlations between hippocampal encoding and recollection effects, associative recognition performance (pR) and baseline memory score, after controlling for age (*N* = 64).

|  | pR |  | baseline memory score |  |
| --- | --- | --- | --- | --- |
|  | *r* | *p* | *r* | *p* |
| *Encoding effect* |  |  |  |  |
| Left hippocampus | **.40** | **.001** | .13 | .324 |
| Right hippocampus | **.38** | **.002** | **.29** | **.024** |
| *Recollection effect* |  |  |  |  |
| Left hippocampus | .24 | .057 | -.09 | .464 |
| Right hippocampus | **.36** | **.004** | -.05 | .682 |

To investigate whether any of the relationships between the hippocampal effects and memory performance were mediated by hippocampal volume, we repeated the foregoing analyses with hippocampal volume as an additional covariate. All of the correlations with pR listed in Table 4 remained significant (partial *r*s > .36, *p*s < .005). However, the relationship between right hippocampal encoding effect and baseline memory scores was not significant after controlling for right hippocampal volume (partial *r* = .24, *p* = .059).

Finally, we examined the direct association between hippocampal volume and pR or baseline memory scores. In contrast to the findings for the functional effects, hippocampal volume was not significantly correlated with pR (for left hippocampus, *r* = .11, *p* = .401; for right hippocampus, *r* = .01, *p* = .922) or baseline memory scores (for left hippocampus, *r* = .02, ***p*** = .852; for right hippocampus, *r* = -.10, *p* = .418).

### 3.5 Predictors of longitudinal memory change

Based on the general model described in the Materials and Methods (see ‘statistical analyses’), four linear mixed effects models (Models 1-4) - one for each of the hippocampal effects (i.e. left hippocampal encoding effect, right hippocampal encoding effect, left hippocampal recollection effect, and right hippocampal recollection effect) - were constructed. For each model, we were interested in: 1) the contribution of the hippocampal effect, which reflects the strength of the relationship between the effect and mean memory performance, and 2) the hippocampal effect × session interaction, which indexes the relationship between the hippocampal effect and memory change.

Results for each model are shown in Table 5. As is evident from the table, the right hippocampal encoding effect (Model 2) was a significant predictor of overall memory performance in the absence of a significant interaction between the effect and session. This finding is consistent with the results reported in Table 4 and Figure 3.

**Table 5.** Linear mixed effects regression results for the encoding- and recollection-related hippocampal effects predicting memory performance and memory change.

| Parameter | B (SE) | df | *t* | *p* |
| --- | --- | --- | --- | --- |
| *Model 1* |  |  |  |  |
| Intercept | 12.07 (6.16) | 50 | 1.96 | .056 |
| Age | -.18 (.09) | 50 | -1.96 | .056 |
| Left_hippo_Enc | .42 (.50) | 58 | .84 | .405 |
| Session | -.15 (.19) | 51 | -.79 | .433 |
| Left_hipp_Enc × Session | -.00 (.27) | 51 | -.01 | .993 |
| *Model 2* |  |  |  |  |
| Intercept | 8.96 (6.07) | 50 | 1.48 | .147 |
| Age | -.13 (.09) | 50 | -1.46 | .150 |
| **Right_hippo_Enc** | **1.18 (.54)** | **58** | **2.18** | **.033** |
| Session | -.15 (.18) | 51 | -.82 | .417 |
| Right_hipp_Enc × Session | .05 (.30) | 51 | .18 | .855 |
| *Model 3* |  |  |  |  |
| **Intercept** | **12.76 (6.33)** | **50** | **2.02** | **.049** |
| Age | -.18 (.09) | 50 | -1.99 | .052 |
| Left_hippo_Re | -.32 (.45) | 56 | -.71 | .479 |
| **Session** | **-.52 (.20)** | **51** | **-2.55** | **.014** |
| **Left_hipp_Re × Session** | **.71 (.22)** | **51** | **3.23** | **.002** |
| *Model 4* |  |  |  |  |
| **Intercept** | **12.48 (6.21)** | **50** | **2.01** | **.050** |
| Age | -.18 (.09) | 50 | -2.00 | .051 |
| Right_hippo_Re | -.09 (.47) | 57 | -.19 | .852 |
| Session | -.33 (.20) | 51 | -1.67 | .102 |
| **Right_hipp_Re × Session** | **.52 (.25)** | **51** | **2.12** | **.040** |

*Note:* Left\_hippo\_Enc: Left hippocampal encoding effect; Right\_hippo\_Enc: Right hippocampal encoding effect; Left\_hipp\_Re: Left hippocampal recollection effect; Right\_hippo\_Re: Right hippocampal recollection effect.

As is also evident from Table 5, in contrast with Model 2, for Models 3 and 4 the hippocampal recollection effects significantly interacted with test session. Since we used baseline scores (the mean of Sessions 1 and 2) as the reference session, these results indicate that the hippocampal recollection effects, especially those in the left hippocampus, were inversely related to longitudinal memory decline. To visualize these effects, we plotted age-residualized hippocampal recollection effects against residualized memory change in Figure 4.

**Figure 4.**
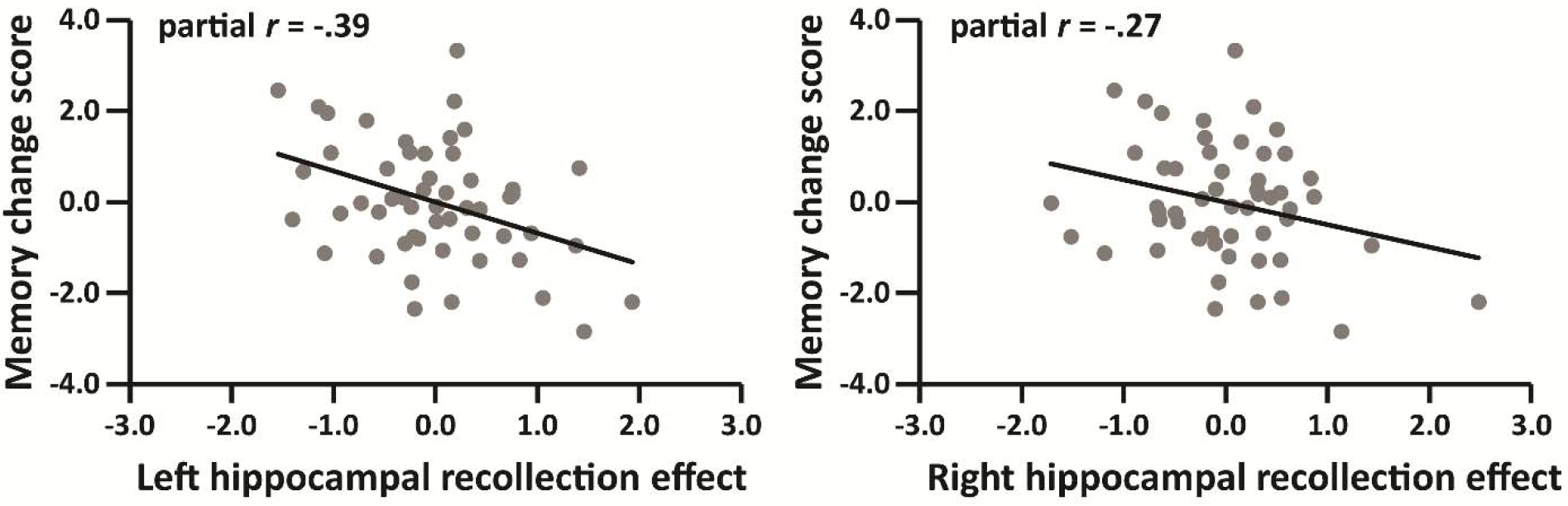
Correlations between memory change scores and left and right hippocampal recollection effects, after controlling for age (*N* = 53).

To examine the possible role of hippocampal volume in mediating these relationships, we constructed follow-up regression models in which either left or right hippocampal volume and the hippocampal volume × session interaction were entered in Models 2, 3 and 4 as additional predictor variables. In the presence of these additional variables, neither the relationship between the right hippocampal encoding effect and memory performance, nor the interaction between the right hippocampal recollection effect and session, were significant [respectively: *t*(56) = 1.57, *p* = .123), *t*(49) = 1.54, *p* = .130]. However, the interaction between the left hippocampal recollection effect and session remained significant [B = .66, *t*(49) = 2.79, *p* = .007].

We also performed two linear mixed effects analyses to directly examine the relationship between hippocampal volume and memory performance or change. The two models included either left or right hippocampal volume, session and hippocampal volume × session as independent variables of interest, and longitudinal memory performance as the dependent variable. We did not identify a significant effect in either model (*p*s > .222).

### 3.6 Correlations among hippocampal functional effects and hippocampal volume

Simple correlations between hippocampal encoding effects, hippocampal recollection effects and hippocampal volume are given in Table 6. As is evident from the table, the encoding effect did not significantly correlate with the ipsilateral recollection effect in either hemisphere. Furthermore, neither the left nor the right hemisphere functional effects correlated with their respective hippocampal volumes.

**Table 6.** Simple correlations among hippocampal encoding effects, hippocampal recollection effects and hippocampal volumes in the full group (*N* = 65 for hippocampal volume, *N* = 64 for hippocampal effects).Right_hippo_Re: Right hippocampal recollection effect; Right_hippo_vol: Right hippocampal volume. Volume measures were residualized against ICV.

| Variable | By variable | *r* | *p* |
| --- | --- | --- | --- |
| *Ipsilateral correlations* |  |  |  |
| Left_hippo_Enc | Left_hippo_Re | -.01 | .966 |
| Left_hippo_vol | Left_hippo_Enc | -.13 | .304 |
| Left_hippo_vol | Left_hippo_Re | -.06 | .656 |
| Right_hippo_Enc | Right_hippo_Re | -.00 | .974 |
| Right_hippo_vol | Right_hippo_Enc | -.15 | .241 |
| Right_hippo_vol | Right_hippo_Re | -.22 | .080 |
| *Contralateral correlations* |  |  |  |
| Left_hippo_Enc | Right_hippo_Enc | **.59** | **< .001** |
| Left_hippo_Re | Right_hippo_Re | **.65** | **< .001** |
| Left_hippo_vol | Right_hippo_vol | **.78** | **< .001** |

*Note.* Left\_hippo\_Enc: Left hippocampal encoding effect; Left\_hippo\_Re: Left hippocampal recollection effect; Left\_hippo\_vol: Left hippocampal volume; Right\_hippo\_Enc: Right hippocampal encoding effect;

As is also evident from Table 6, in contrast to the ipsilateral correlations, there were robust positive across-hemisphere correlations for both classes of functional effect. Similarly, left and right hippocampal volumes were positively correlated.

## 4 Discussion

The primary aim of the present study was to investigate the relationships between encoding- and recollection-related hippocampal effects, memory performance and three-year longitudinal memory change in a sample of healthy older adults. We found that while right hippocampal encoding effects were correlated with baseline memory performance, both left and right hippocampal recollection effects were predictive of memory change. To our knowledge, this is the first evidence to link hippocampal recollection effects obtained at baseline to longitudinal memory change in older adults.

Before discussing these findings, we note that re-test effects can pose significant obstacles for the interpretation of longitudinal data. Such effects can persist over several years and lead to the underestimation of cognitive change (Nyberg et al., 2016; Salthouse, 2009). In light of prior findings that re-test effects for verbal memory tend to diminish after the first re-test session (Salthouse and Tucker, 2008), we adopted a burst measurement design (Salthouse and Nesselroade, 2010) and re-administered the neuropsychological test battery shortly after the first test session. For the reasons outlined in the Materials and Methods, we elected to employ test scores averaged over these initial two sessions to estimate baseline performance. When assessed against this baseline, memory performance did not demonstrate a reliable decline over the follow-up period at the group level. However, this finding should not necessarily be taken as evidence that the memory performance of our sample remained stable over this period. Notably, if it is assumed that session 2 performance provides the best correction for session 3 re-test effects, then memory scores declined significantly and robustly at the group level (see Table 3).

Given that there is no basis for preferring one of these measures of baseline performance over the other (or any other weighting of session 1 and session 2 performance), we elected to employ arguably the most stable measure, and to interpret the ensuing metric of memory change in relative rather than absolute terms. That being said, as is reported in the Supplementary Material, our main findings were unaffected when session 2 scores alone were employed as the baseline measure.

As noted in the Introduction, prior fMRI studies examining across-participants relationships between encoding- and recollection-related hippocampal effects and memory performance focused on measures of performance derived from the memory task employed to estimate the fMRI effects (Daselaar et al., 2006; de Chastelaine et al., 2016a, 2016b; Wang et al., 2016). The novel aspect of the present study is the extension of these analyses to ‘offline’ neuropsychological measures of baseline memory performance and its change over time. However, it should be noted that the positive age-invariant correlations between hippocampal encoding and recollection effects with in-scanner memory performance described in our prior reports (de Chastelaine et al., 2016a, 2016b) were also evident in the present sample, which comprised a subgroup of the 136 participants described in those reports. The present findings lend credence to the proposal that these fMRI effects index the efficacy of functionally significant mnemonic processes in older adults. Indeed, a multiple regression model predicting the in-scanner associative recognition performance of our baseline sample of 64 older participants from their four hippocampal functional effects (i.e. left and right encoding and recollection effects) accounted for more than 30% of the variability in performance (adjusted *R* = .318, *p* < .001).

As already noted, the principal focus of the present study was not on the relationship between fMRI effects and in-scanner memory performance, but rather, the relationship between these effects and memory metrics derived from standardized neuropsychological test scores. In the case of baseline performance, we identified a significant positive relationship between baseline scores and the right hippocampal encoding effect, and obtained a convergent result from the linear mixed effects model (Model 2) that employed this fMRI effect as a predictor of memory performance in the longitudinal subgroup (although, obviously, these two findings should not be viewed as independent). Turning to the longitudinal component of the study, we found that both hippocampal recollection effects were predictive of memory change, although the relationship in the left hippocampus was the more robust.

Whereas the sizeable correlations between in-scanner memory performance and hippocampal encoding and recollection effects point to the functional significance of both classes of effect, this does not mean that they reflect common, or even closely related, cognitive operations. Moreover, the finding that the across-subject correlations between two classes of effect were essentially zero (Table 6) indicates that the effects do not both reflect individual differences in some ‘trait-like’ factor such as hippocampal functional integrity or efficacy. Arguably, these findings are understandable given the differing roles proposed for the hippocampus during encoding and retrieval (e.g. Rugg et al., 2016). At encoding, the hippocampus is held to be responsible for ‘binding’ patterns of cortical activity elicited by an event into a sparse, content-addressable memory representation. As was discussed in de Chastelaine et al. (2016a), in light of this proposed role, the relationship between hippocampal encoding effects and memory performance could be an indirect rather than a direct one. That is, the relationship might reflect individual differences, not in the functional efficacy of the hippocampus, but in the amount or the quality of the information about a study event that it receives. For example, there is evidence that both subsequent memory performance and hippocampal encoding effects are sensitive to the amount of attentional resources that are directed toward a study event or a subset of its features (e.g. Aly and Turk-Browne, 2016; Uncapher and Rugg, 2009). Therefore, the present findings for associative memory performance might be a reflection of individual differences in the processing resources or attentional strategies engaged by participants while they performed the study task. The finding that (right) hippocampal encoding effects also predicted baseline memory performance suggests that these individual differences also contributed to across participant variability in baseline memory scores. The additional finding that pR and baseline memory performance were robustly correlated adds credence to this proposal.

The contribution of the hippocampus to successful episodic retrieval is distinct from that at encoding. Recollection is held to occur when a retrieval cue activates a hippocampal memory representation sufficiently to give rise to ‘pattern completion’, which restores the representation to an active state. In turn, this leads to reinstatement of the encoded pattern of cortical activity, providing access to mnemonic content (see Rugg et al., 2016, for review). From this perspective, therefore, the determinants of the magnitude of hippocampal recollection effects are distinct from those moderating encoding effects, and will include such factors as the efficacy of cue processing and the amount and specificity of the information retrieved in response to the cue (Mayes et al., 2019; Rugg et al., 2012). Since these factors would be expected to contribute to memory performance, it is perhaps unsurprising that hippocampal recollection effects correlated significantly with associative recognition performance in the present study, while at the same time correlating negligibly with hippocampal encoding effects.

Why was memory change correlated with hippocampal recollection effects, but not with encoding effects? We conjecture that this dissociation reflects a combination of two factors. First, that age-related memory change largely reflects a decline in the ability to recollect qualitative information about past events, rather than in memory processes that do not depend heavily on the hippocampus, such as familiarity (see Koen and Yonelinas, 2014, for review of the extensive cross-sectional literature supporting this contention). Second, that hippocampal recollection effects provide a ‘purer’ or more direct index of the structure’s contribution to recollection than do encoding effects which, as noted previously, are likely sensitive to a multiplicity of processes that depend on extra-hippocampal regions. The relationship between hippocampal recollection effects and memory change can then be explained if it is assumed that the effects are indicative of both the current functional integrity of the structure, *and* its capacity to resist or cope with future age-related functional degradation. Establishing the validity of this account seems a worthy goal for future research.

In the present study, all correlations between functional hippocampal effects and memory performance were positive: larger effects predicted higher performance on the in-scanner memory task, and higher baseline memory scores and less decline in these scores over three years. These findings stand in contrast to those from other studies where either a null (Dulas and Duarte., 2011, 2016) or a negative relationship (Carr et al., 2017; Daselaar et al., 2015; Miller at al., 2008) between hippocampal memory effects and memory performance was reported. Whereas null findings can plausibly be attributed to any number of factors that might have obscured a ‘true’ relationship, notably, lack of statistical power arising from small sample sizes, findings of a negative relationship clearly conflict with the present results (and those of, for example, Daselaar et al., 2005; de Chastelaine et al., 2016a, 2016b; and Wang et al., 2016). One possibility is that these disparate findings reflect variation in the cognitive status of the older adult samples employed in the different studies. Notably, it has been reported that older adults with mild cognitive impairment (MCI) likely attributable to prodromal Alzheimer’s Disease (AD) demonstrate hippocampal ‘hyperactivity’ - an elevation of task-related hippocampal responses relative to age-matched, low-risk controls (e.g. Bakker et al., 2015; Dickerson et al., 2004; Putcha et al., 2011). Moreover, negative correlations between task-related hippocampal activity and memory performance have been reported in MCI samples (Bakker et al., 2012; Yassa et al., 2010). Thus, if samples of older adults include a sufficient number of individuals at high risk for AD, a negative relationship between hippocampal memory effects and performance might be anticipated. In the present study, participants were screened to exclude individuals with cognitive profiles or medical histories indicative of elevated risk for prodromal AD (see Materials and Methods). Since no participant in our longitudinal subgroup transitioned to MCI status over the three-year follow-up period, we assume the screening procedure was reasonably effective. A similar approach was adopted in Miller et al. (2008), but it is perhaps noteworthy that the participants included in the study of Carr et al. (2017) comprised a mixture of healthy and cognitively impaired older adults. No information about the cognitive profiles of the participants employed in Daselaar et al. (2015) was provided in that report.

In contrast to the functional effects, we were unable to identify significant relationships between hippocampal volume and either baseline memory performance or memory change. Perhaps unsurprisingly in light of these null findings, hippocampal volume also had little or no moderating influence on the relationship between the functional effects and these behavioral measures. The present null findings in respect of hippocampal volume are not without precedent. The numerous prior studies examining the relationship between hippocampal volume and memory performance in healthy older adults have yielded an inconsistent pattern. Whereas some studies reported a positive correlation (e.g. Ezzati et al., 2016; O’Shea et al., 2016; Rosen et al., 2003), others have failed to find such evidence (e.g. Charlton et al., 2010; Walhovd et al., 2010; for review, see Kaup et al., 2011; Van Petten, 2004). Similar inconsistencies also exist for longitudinal studies examining relationships between hippocampal volume and memory decline (see Gorbach et al., 2017; Mungas et al., 2005 for examples of positive findings; see Cardenas et al., 2011; Carmichael et al., 2012 for examples of null results; for review, see Oschwald et al., in press).

Finally, we note three limitations of the present study. First, the sample size was modest, limiting statistical power and constraining the size of the effects that could be detected. Second, since we only assessed memory performance on what was, effectively, two occasions, we were unable to characterize the trajectory of memory change in our participants. This limitation is compounded by the relatively short follow-up period of three years. Third, the associations with hippocampal effects identified here could conceivably reflect individual differences not only in neural activity but in one or more vascular factors, such as cerebrovascular reactivity (CVR) - an important non-neural determinant of BOLD signal magnitude (e.g. Liu et al., 2013; Lu et al., 2011; Tsvetanov et al., 2015). Since we did not control for CVR, we cannot rule out some influence of this variable. Clearly, future research would benefit from the employment of larger samples subjected to multiple test sessions over a longer overall follow-up period, along with functional methods that correct for or which are insensitive to individual differences in neurovascular coupling.

These limitations notwithstanding, consistent with prior findings reviewed in the Introduction, the present results suggest that hippocampal functional activity is predictive of both individual differences in memory performance and longitudinal memory change in cognitively unimpaired older adults. Going beyond prior reports, the results further suggest that experimental contrasts that isolate the role of the hippocampus in recollection-based memory judgments might hold promise as predictors of future memory performance in this population.

## Acknowledgements

We acknowledge the contributions of Hannah Stanton and Kay Moolenijzer for their assistance with participant recruitment and neuropsychological data collection. We also thank the staff of the UTSW Advanced Imaging Center for their assistance with MRI data collection.

