## supplemental materials for "Recollection-related hippocampal fMRI effects predict longitudinal memory change in healthy older adults"

**Supplementary Material**

1. Rotated factor loadings from the PCA of the neuropsychological test data

2. Functional hippocampal effects in the longitudinal subgroup

3. Correlations between functional and structural measures and memory performance in the longitudinal subgroup

4. Correlations among hippocampal functional effects and hippocampal volume in the longitudinal subgroup

5. Predictors of session 2 memory performance

6. Predictors of longitudinal change with session 2 scores indexing baseline memory performance

***1. Rotated factor loadings from the PCA of the neuropsychological test data***

The PCA conducted on the session 1 test data (see Materials and Methods) yielded a 4-component solution which explained 66.8% of the variance in the data. As can be seen from Supplemental Table 1, after rotation the components represented the cognitive constructs (broadly defined) of memory (C1), fluency (C2), speed (C3) and crystallized intelligence (C4).

Supplemental Table 1. Rotated factor loadings from the PCA of the neuropsychological test data from older adults included in Session 1 (*N* = 67).

| Task | Component | | | |
| --- | --- | --- | --- | --- |
|  | Memory  (C1) | Fluency  (C2) | Speed  (C3) | Crystallized IQ (C4) |
| CVLT Hits | .68 | -.31 | .05 | .37 |
| CVLT False Alarms | -.54 | -.25 | .27 | .41 |
| CVLT Composite | .86 | .08 | -.17 | -.02 |
| FAS | .07 | .68 | .01 | .42 |
| Logical Memory Composite | .67 | .05 | -.06 | .13 |
| SDMT | .30 | .65 | -.36 | .02 |
| Trail A | -.05 | -.19 | .89 | -.10 |
| Trail B | -.12 | -.08 | .86 | -.30 |
| Digit Span | .11 | .11 | -.16 | .79 |
| Category Fluency (Animals) | -.04 | .78 | -.14 | .04 |
| WTAR (Full-Scale IQ) | .09 | .20 | -.30 | .70 |
| Raven’s | .60 | .43 | .05 | .02 |

***2. Functional hippocampal effects in the longitudinal subgroup***

We examined whether the functional effects were reliable at the group level, and whether there was any evidence of lateralization in the effects. For the study data, we conducted a 2 (Condition: later associative hit vs. later associative miss) × 2 (Hemisphere: left vs. right) on the parameter estimates for later associative hits and misses extracted from the hippocampal ROIs. Neither the main effect of condition nor the condition × hemisphere interaction was significant, *p*s > .086, partial *η*^2^s < .06. An analogous ANOVA conducted on the parameter estimates for associative hits and misses at retrieval revealed a significant effect of condition, *F*(1, 52) = 20.92, *p* < .001, partial *η*^2^ = .29, indicative of greater hippocampal activity for associative hits (*M* = -.02) than for associative misses (*M* = -.45). This effect interacted significantly with hemisphere, *F*(1, 52) = 4.30, *p* = .043, partial *η*^2^ = .08, reflecting larger recollection effects in the left hemisphere. Simple effects analyses indicated that recollection effects were reliable in both hemispheres (*p*s < .002).

***3. Correlations between functional and structural measures and memory performance in the longitudinal subgroup***

Correlations between hippocampal encoding effects, hippocampal recollection effects and baseline pR for the longitudinal subgroup are shown in Supplemental Table 2. As is evident from this table, with only one exception, these effects were significantly positively correlated with pR.

Supplemental Table 2. Correlations between encoding- and recollection-related hippocampal effects and pR and baseline memory score, after controlling for age (*N =* 53).

|  | pR | | baseline memory score | |
| --- | --- | --- | --- | --- |
|  | *r* | *p* | *r* | *p* |
| *Encoding effect* |  |  |  |  |
| Left hippocampus | .45 | **.001** | .13 | .365 |
| Right hippocampus | .33 | **.018** | .32 | **.021** |
| *Recollection effect* |  |  |  |  |
| Left hippocampus | .15 | .300 | -.10 | .483 |
| Right hippocampus | .29 | **.040** | -.02 | .870 |

To investigate whether any relationships between the hippocampal effects and either pR or baseline memory scores were mediated by hippocampal volume, we repeated the foregoing analyses after adding hippocampal volume as an additional covariate. All of the relationships with pR identified above remained significant (partial *r*s > .30, *p*s < .030). The relationship between right hippocampal encoding effect and baseline memory score was however non-significant, partial *r* = .25, *p* = .086.

We also examined the direct association between hippocampal volume and baseline pR. In contrast to the findings for the functional effects, hippocampal volume was not significantly correlated with pR (for left hippocampus, *r* = .11, *p* = .444; for right hippocampus, *r* = .01, *p* = .934). Similarly, hippocampal volume was not significantly correlated with baseline memory scores (for left hippocampus, *r* = -.02, *p* = .885; for right hippocampus, *r* = -.14, *p* = .321).

***4. Correlations among hippocampal functional effects and hippocampal volume in the longitudinal subgroup***

Simple correlations between hippocampal encoding effects, hippocampal recollection effects and hippocampal volume in the longitudinal subgroup are given in supplemental Table 3. As is evident from the table, most ipsilateral correlations were not significant, with the exception that right hippocampal volume was negatively correlated with the right hippocampal encoding effect. In contrast to the ipsilateral correlations, there were robust positive across-hemisphere correlations for both classes of functional effect. Left and right hippocampal volumes were also positively correlated.

Supplemental Table 3. Simple correlations among hippocampal encoding effect, hippocampal recollection effect and hippocampal volume in the full group (*N* = 54 for hippocampal volume, *N* = 53 for hippocampal effects).

| Variable | By variable | *r* | *p* |
| --- | --- | --- | --- |
| *Ipsilateral correlations* | | | |
| Left_hippo_Enc | Left_hippo_Re | .05 | .745 |
| Left_hippo_volume | Left_hippo_Enc | -.24 | .089 |
| Left_hippo_volume | Left_hippo_Re | -.01 | .958 |
| Right_hippo_Enc | Right_hippo_Re | -.05 | .710 |
| Right_hippo_volume | Right_hippo_Enc | **-.32** | **.022** |
| Right_hippo_volume | Right_hippo_Re | -.19 | .173 |
| *Contralateral correlations* | | | |
| Left_hippo_Enc | Right_hippo_Enc | **.61** | **< .001** |
| Left_hippo_Re | Right_hippo_Re | **.65** | **< .001** |
| Left_hippo_volume | Right_hippo_volume | **.79** | **< .001** |

*Note*. Left_hippo_Enc: Left hippocampal encoding effect; Right_hippo_Enc: Right hippocampal encoding effect; Left_hipp_Re: Left hippocampal recollection effect; Right_hippo_Re: Right hippocampal recollection effect; Left_hippo_volume: Left hippocampal volume; Right_hippo_volume: Right hippocampal volume. Volume measures were residualized against ICV.

***5. Predictors of session 2 memory performance***

Controlling for age, we repeated the partial correlation analyses to examine the relationship between recollection-related hippocampal effects and baseline memory indexed by session 2 scores in the full group. Similar to the results reported in the main text, only the right hippocampal encoding effect was significantly correlated with session 2 scores, partial *r* = .26, *p* = .04 (for the rest of the partial correlations, absolute partial *r*s < .13, *p*s > .350). This relationship did not survive when right hippocampal volume was added as an additional covariate (partial *r* = .23, *p* = .081).

The same pattern was observed in the longitudinal subgroup: The right hippocampal encoding effect, but not the other hippocampal effects (absolute partial *r*s < .14, *p*s > .323), was significantly correlated with session 2 scores (partial *r* = .31, *p* = .026). However, when right hippocampal volume was controlled for, the relationship between right hippocampal encoding effect and session 2 scores was non-significant (partial *r* = .25, *p* = .077).

Also consistent with the results reported in the main text, hippocampal volumes were not significantly correlated with the session 2 memory component scores (for left hippocampus, *r* = .04, *p* = .730; for right hippocampus, *r* = -.10, *p* = .426) in the full group. Similarly, we did not observe significant correlations between hippocampal volume and session 2 scores in the longitudinal subgroup (for left hippocampus, *r* = .01, *p* = .939; for right hippocampus, *r* = -.13, *p* = .334).

***6. Predictors of longitudinal change with session 2 scores indexing baseline memory performance***

Similarly to what we describe in the main text, we constructed four linear mixed models to investigate relationships between hippocampal effects and mean memory performance and longitudinal memory change from session 2 to session 3.

Results for each model are shown in Supplemental Table 4. Consistent with results reported for the group-level comparison in the main text (see Table 3), in all models there was a significant main effect of session, indicative of a significant memory decline from session 2 to session 3.

As is also evident from Supplemental Table 4, the right hippocampal encoding effect (Model 2) was a significant predictor of overall memory performance. By contrast, the left hippocampal recollection effect (Model 3) significantly interacted with test session. The interaction between right hippocampal recollection effect and test session did not achieve significance (Model 4).

To examine the possible role of hippocampal volume in mediating these relationships, we constructed follow-up regression models in which either left or right hippocampal volume and the hippocampal volume × session interaction were entered in Models 2 and 3 as additional predictor variables. In the presence of these additional variables, the relationship between the right hippocampal encoding effect and memory performance was not significant, *t*(56) = 1.61, *p* = .113. However, the interaction between the left hippocampal recollection effect and session remained significant [B = .75, *t*(49) = 3.35, *p* = .002].

We also performed two linear mixed effects analyses to directly examine the relationship between hippocampal volume and memory performance or change. The two models included either left or right hippocampal volume, session and hippocampal volume × session as independent variables of interest, and longitudinal memory performance as the dependent variable. Other than a significant main effect of session in each model (*p*s < .001), we did not observe any significant effect of hippocampal volume or hippocampal volume × session in either model (*p*s > .381).

Supplemental Table 4. Linear mixed effects regression results for recollection-related hippocampal effects predicting memory performance and memory change.

| Parameter | B (SE) | df | *t* | *p* |
| --- | --- | --- | --- | --- |
| *Model 1* | | | | |
| **Intercept** | **13.04 (6.06)** | **50** | **2.15** | **.036** |
| **Age** | **-.18 (.09)** | **50** | **-2.03** | **.048** |
| Left_hippo_Enc | .45 (.49) | 58 | .90 | .371 |
| Session | -.91 (.18) | 51 | -4.97 | **<.001** |
| Left_hipp_Enc × Session | -.03 (.26) | 51 | -.11 | .916 |
| *Model 2* | | | | |
| Intercept | 10.08 (5.99) | 50 | 1.68 | .099 |
| Age | -.13 (.09) | 50 | -1.54 | .129 |
| **Right_hippo_Enc** | **1.11 (.53)** | **58** | **2.08** | **.043** |
| **Session** | **-.92 (.18)** | **51** | **-5.14** | **<.001** |
| Right_hipp_Enc × Session | .12 (.29) | 51 | .42 | .677 |
| *Model 3* | | | | |
| **Intercept** | **13.81 (6.23)** | **50** | **2.22** | **.031** |
| **Age** | **-.19 (.09)** | **50** | **-2.07** | **.044** |
| Left_hippo_Re | -.35 (.44) | 56 | -.79 | .432 |
| **Session** | **-1.30 (.19)** | **51** | **-6.70** | **<.001** |
| **Left_hipp_Re × Session** | **.74 (.21)** | **51** | **3.52** | **.001** |
| *Model 4* | | | | |
| **Intercept** | **13.41 (6.11)** | **50** | **2.20** | **.033** |
| **Age** | **-.18 (.09)** | **50** | **-2.06** | **.045** |
| Right_hippo_Re | -.02 (.47) | 57 | -.05 | .961 |
| **Session** | **-1.07 (.19)** | **51** | **-5.60** | **<.001** |
| Right_hipp_Re × Session | .45 (.24) | 51 | 1.89 | .064 |

*Note*: Left_hippo_Enc: Left hippocampal encoding effect; Right_hippo_Enc: Right hippocampal encoding effect; Left_hipp_Re: Left hippocampal recollection effect; Right_hippo_Re: Right hippocampal recollection effect.
